# A Multiplex Droplet Digital PCR Assay for Chromosome Copy Number Determination in *Candida albicans*

**DOI:** 10.64898/2026.02.09.704919

**Authors:** Eli Isael Maciel, Sylvain Ursuegui, Sana Ahmed-Seghir, Corinne Maufrais, Stéphanie Roy, Cecile Gautier, Philipp Brandt, Cécile Jovelet, Laras Pitayu, Iuliana V. Ene

## Abstract

Chromosome copy number variation (CNV) is a major contributor to genome plasticity and adaptation in *Candida albicans*, a leading fungal pathogen of humans. Aneuploidy, defined as deviations from the normal diploid chromosome set, rapidly alters gene dosage, enabling tolerance to host-imposed and antifungal stress. Accurate detection and quantification of chromosomal copy number changes are thus essential to dissect the mechanisms by which *C. albicans* adapts and evolves. Here, we describe the development, optimization, and validation of a six-color, 16-plex droplet digital PCR assay for simultaneous quantification of all *C. albicans* chromosome arms in a single reaction. Each target is detected by a unique dual-color or single-color combination of probes, enabling high-order multiplexing through binary fluorescence encoding. Following optimization of probe concentrations, PCR cycling parameters, genomic DNA extraction and pre-treatment with restriction enzymes, the assay provides accurate, reproducible chromosome-level copy number estimates that correlate closely with WGS results across euploid and aneuploid isolates. Compared to whole-genome sequencing, the assay is rapid, cost-effective, and scalable, requiring minimal DNA input and allowing high-throughput analysis of large isolate collections. The 16-plex assay thus provides a platform for dissecting genome instability and adaptive evolution in *C. albicans*.

**Article Summary:** We developed and validated a 16-plex droplet digital PCR assay that estimates chromosome dosage across the entire genome of the human fungal pathogen *C. albicans* in a single reaction. The assay uses six fluorescent colors and unique color combinations to track one marker on each chromosome arm, enabling rapid detection of aneuploidy (extra or missing chromosomes). Results closely matched whole-genome sequencing for isolates with simple aneuploid forms and detected low-frequency trisomic clones in mixed populations. With optimized DNA preparation, this method provides a practical tool for screening genome instability in research and clinical settings.

## Introduction

Aneuploidy, the presence of abnormal numbers of chromosomes, is a hallmark of genome instability and a major driver of adaptive evolution across diverse biological systems (Torres et al. 2008; Li and Zhu 2022). In cancer, aneuploidy fuels tumor heterogeneity, promotes rapid adaptation to therapy, and contributes to disease progression (Li and Zhu 2022; Vasudevan et al. 2021). The capacity for karyotypic plasticity is particularly evident in human fungal pathogens, where aneuploidy has emerged as a pervasive mechanism of adaptation to host environments and antifungal stress. Here, it enables rapid adaptation to environmental and drug-induced stress by altering gene dosage without requiring point mutations (Todd and Selmecki 2020; Tsai and Nelliat 2019; Bennett et al. 2014; Vande Zande et al. 2023).

Among fungal pathogens, *C. albicans* exemplifies the dual nature of aneuploidy as both a source of genetic innovation and a contributor to phenotypic variability. *C. albicans* is an opportunistic commensal capable of causing life-threatening systemic infections in immunocompromised individuals (Denning 2024; Brown et al. 2012; Pappas et al. 2018). The remarkable genomic plasticity of this species underlies its ability to thrive in fluctuating host niches and in the presence of antifungal drugs (Selmecki et al. 2010). Unlike most fungi, *C. albicans* lacks a complete sexual cycle, limiting its ability to shuffle alleles through meiosis (Ene and Bennett 2014). Instead, it exploits single point mutations, aneuploidy, loss of heterozygosity, and parasexual recombination to generate genetic diversity (Vande Zande et al. 2023; Todd et al. 2019; Forche 2014; Morrow and Fraser 2013; Selmecki et al. 2010; Bennett et al. 2014).

Aneuploidy has been repeatedly observed in clinical isolates and experimental populations, often associated with adaptation to host niches and antifungal resistance. For example, monosomy of chromosome (Chr) 5 promotes tolerance to fluconazole and caspofungin (Yang et al. 2019), trisomy of Chr2 confers cross-resistance to caspofungin, tunicamycin, and hydroxyurea (Yang et al. 2021a), while duplication of the left arm of Chr5 (iso-Chr5L) drives fluconazole resistance (Selmecki et al. 2008; Selmecki et al. 2006). Despite frequent fitness trade-offs associated with aneuploidy (Zhu et al. 2018; Yang et al. 2021b; Tsai et al. 2019; Tsai and Nelliat 2019; Torres et al. 2007), aneuploid isolates can also facilitate adaptation to mammalian host niches. Acquisition of a Chr7 trisomy enhances colonization of the murine gastrointestinal tract by increasing *NRG1* copy numbers and suppressing filamentation (Ene et al. 2018; Kakade et al. 2023). Trisomy of Chr5 and Chr6 increases fitness for the murine oral cavity, which was associated with decreased epithelial adherence and invasion (Forche et al. 2018). These findings highlight aneuploidy as a powerful, reversible adaptive mechanism that enables *C. albicans* to fine-tune its physiology in response to environmental and host-imposed stresses, balancing the costs of genomic instability with context-dependent fitness gains that enhance survival, persistence, and pathogenic potential.

Accurate determination of chromosome copy number and identity is therefore critical for dissecting the role of aneuploidy in fungal adaptation. Whole-genome sequencing (WGS) remains the gold standard for this purpose, providing genome-wide resolution and quantitative accuracy. However, its cost, turnaround time, and computational requirements limit its use in large-scale or high-throughput studies. Alternative methods, such as flow cytometry and quantitative PCR (qPCR), provide faster and more accessible options but suffer from limitations: flow cytometry detects total DNA content without chromosome identity, while qPCR is constrained by limited multiplex capacity and reliance on reference gene normalization.

Droplet digital PCR (ddPCR) enables accurate absolute quantification by partitioning a sample into thousands of nanodroplets in which amplification proceeds independently. Applying Poisson statistics to the fraction of positive droplets yields precise copy-number estimates without the need for standard curves. In conventional digital PCR (dPCR), each target is detected with a specific fluorescent probe in a single detection channel, so multiplexing capacity is constrained by the limited number of optical channels. To overcome this, higher-order multiplexing methods use different fluorescence intensities (amplitude- or ratio-based approaches) to encode multiple targets within the same channel. However, these methods require complex multi-level thresholding and are less robust because fluorescence intensity can vary across samples. An alternative approach is color-combination, in which a unique color signature defines each target. While this is impractical in qPCR (because overlapping bulk fluorescence signals create ambiguity), dPCR can precisely identify color combinations in each partition and exclude ambiguous co-encapsulation events. This color-combination approach has been successfully implemented on six-color and seven-color multiplex dPCR platforms, enabling detection of up to 15 and 21 targets, respectively. Importantly, the analysis remains simple and robust: only one threshold per fluorescence channel is required, and dedicated analysis software can automatically identify and quantify targets based on predefined color codes (Santos-Barriopedro 2023). This approach substantially expands multiplexing capacity beyond conventional methods and minimizes competition between primer-probe sets.

In this study, we designed, optimized, and validated a 16-plex ddPCR assay for *C. albicans* that measures copy number across all 16 chromosome arms in a single reaction. The panel enables genome-wide quantification with single-reaction throughput and accuracy comparable to WGS. We describe the assay development process, including probe design and refinement of underperforming loci, optimization of DNA extraction and digestion, PCR cycling conditions, and calibration of fluorescence compensation. We then show that the assay detects mixed euploid-aneuploid populations and validate its quantitative precision by comparison with WGS-derived copy number profiles across a diverse strain panel. Together, these results establish the 16-plex ddPCR assay as a rapid, scalable, and quantitative approach for genome-wide aneuploidy detection in *C. albicans*.

## Methods

### *C. albicans* strains and growth

All experiments were performed using *C. albicans* isolates representing a range of karyotypes. The reference strain SC5314 (euploid A) served as one of the main euploid controls. For sensitivity testing in mixed populations, disomic and aneuploid subpopulations derived from the same genetic background (euploid B, CAY6440) were combined in defined ratios (25%, 50%, and 75% aneuploid). The collection of isolates used throughout the study is available in **Supplementary Table 1.** Yeast cells were cultured in YPD medium (1% yeast extract, 2% peptone, and 2% dextrose) overnight at 30°C with shaking (200 rpm). Cells were collected from the stationary phase for genomic DNA extraction.

### Genomic DNA extraction

The yeast strains were grown on agar or liquid YPD medium for DNA extraction. Two extraction methods were evaluated for their effects on assay performance: a phenol–chloroform method and a QIAGEN Genomic-tip column-based kit (10223, QIAGEN).

For phenol-chloroform extractions, 2 mL of an overnight culture or a patch of cells from a plate was washed twice with sterile water and resuspended in 400 µL TE (Tris-EDTA, pH 8) buffer containing 20 µL 20% SDS and 200 µL phenol:chloroform:isoamyl alcohol (25:24:1, neutral pH). Cells were lysed by bead beating (FastPrep, three cycles of 20 s at power setting 4, with a 3 min rest on ice between cycles). The aqueous phase was recovered, and the DNA was precipitated with ethanol and washed with 70% ethanol. Residual RNA was removed with RNase treatment.

Column-based extractions were performed according to the manufacturer’s instructions (QIAGEN). Briefly, 2 mL of an overnight culture or a patch of cells from a plate was washed with 1X TE buffer, the cell wall was digested with Lyticase (L2524, Sigma) in buffer Y1 for 1 h, followed by treatment with Proteinase K (P2308, Sigma) and RNase (R5503, Sigma) in buffer G1. The DNA was purified using Genomic-tip columns, eluted, precipitated with isopropanol, washed with 70% cold ethanol, and resuspended in sterile water.

### DNA digestion

Restriction enzyme digestion was tested using FastDigest enzymes (Thermo Fisher) BamHI (FD0054), EcoRI (FD0274), HindIII (FD0504), and PstI (FD0614). Each reaction contained 3 µg of genomic DNA, which was digested for 30 min according to the manufacturer’s recommended conditions. Digested DNA was quantified and diluted to final template concentrations of 10 or 30 pg per reaction. Single-enzyme and combination digestions were tested systematically. SnapGene Viewer was used to verify enzyme restriction sites.

### Target Selection and Probe Design

The *C. albicans* Crystal digital PCR assay (R52002, Stilla Technologies, Bio-Rad Laboratories) was designed to quantify chromosome copy number at 16 loci, one per chromosome arm. Additional assay details, including recommended reaction concentrations, validated restriction enzymes, thermocycling programs, and instructions for image and data analysis, are provided in the assay information sheet (https://go.stillatechnologies.com/assay/c-albicans-crystal-digital-pcr-r-assay).

To minimize sequence or copy number variation that might affect probe binding, candidate loci were chosen based on three principles: near diploid coverage in the SC5314 reference strain, proximity to centromeric regions, and the potential for high gene conservation due to essential or housekeeping function. Each target was verified against the SC5314 reference genome (assembly 22) to confirm the absence of restriction enzyme cut sites within the amplicon (**Supplementary Table 2**). The final targets, fluorophore combinations, and encoding schemes are listed in **Supplementary Table 3**. Probes used in this study were designed using Flex Probe and Universal Reporter Technology (Stilla Technologies, Bio-Rad Laboratories). Flex Probes are target-specific, non-fluorescent probe oligonucleotides that trigger the fluorescence of Universal Reporters in the presence of the targeted sequence. To design the oligonucleotides, the SC5314 assembly 22 was used. Sequence homology with species closely related to *C. albicans* (e.g., *Candida dubliniensis*, *Candida tropicalis*) was assessed using BLAST, and the primer and probe design were adjusted accordingly. Each target was first tested individually (simplex) and then as part of the 16-plex mixture. Simplex reactions yielded higher apparent copy numbers (reflecting reduced competition), but multiplex reactions preserved expected chromosome ratios and produced reproducible results after probe re-design and digestion.

### Reaction mix preparation and PCR cycling conditions

For each reaction, 0.6 µL Crystal Flex probes and primers were mixed with 0.6 µL 10X Buffer A, 0.24 µL 10X Buffer B, 0.15 µL Universal Reporter A, 0.15 µL Universal Reporter B (all naica PCR MIX reagents (R10106) and Universal Reporters 7 (R42401), Stilla Technologies, Bio-Rad Laboratories), and 4.3 µL of 14 or 42 pg/µL genomic DNA (gDNA) template. Primers and Flex Probes are listed in **Supplementary Table 4**. A total of 5 µL of the prepared reaction mixture was loaded into each chamber of a Ruby Chip (C16011, Stilla Technologies, Bio-Rad Laboratories). The chip was loaded into the dPCR cycler (naica Geode, Stilla Technologies, Bio-Rad Laboratories), a device that combines droplet generation and thermocycling. Following droplet partitioning, the amplification protocol was initiated, consisting of an initial denaturation at 95°C for 3 min, followed by 60 cycles of 15 s at 95°C for denaturation and 45 s at 60°C for annealing and elongation. A final elongation step at 58°C for 15 min was performed before chip release. Alternate programs tested included extended elongation (60 s) and final elongation durations (10 or 15 min). Longer elongation at 60 s and a final elongation of 15 min improved separability scores, simplifying thresholding and improving left/right arm balance, especially for low-amplitude targets (e.g., Chr4R). Increasing probe concentrations from 0.5 µM to 1 µM further improved amplitude and reduced uncertainty.

To quantify the discrimination between positive and negative ddPCR droplets, we used a separability score (S), calculated by Crystal Miner, that incorporates both the magnitude of signal separation and the variability within each droplet population (**Supplementary Figure 1**). For each ddPCR reaction, droplets were partitioned into positive and negative populations based on fluorescence amplitude. The separability score was then calculated as:

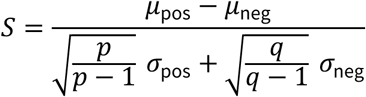

Where *μ*_pos_ and *μ*_neg_ are the mean fluorescence amplitudes of the positive and negative droplet populations, respectively; *σ*_pos_ and *σ*_neg_ are the corresponding standard deviations; *p* and *q* are the number of positive and negative droplets, respectively. A higher score indicates greater separation between positive and negative droplets, reflecting improved resolution. This metric provides a normalized measure of ddPCR performance that accounts for both signal dispersion and population size.

### Fluorescence Detection and Compensation

The chips were scanned using the dPCR reader (naica Prism6, Stilla Technologies, Bio-Rad Laboratories), which is equipped with six fluorescence channels (blue, teal, green, yellow, red, and infrared (IR)). Exposure times were optimized for each channel: Blue = 500 ms, Teal = 800 ms, Green = 250 ms, Yellow = 350 ms, Red = 1000 ms, IR = 1000 ms. To correct for spectral overlap, a spillover compensation matrix (Madic et al. 2016) was generated using mono-color control reactions and implemented in the Crystal Miner v4.0 software (Stilla Technologies, Bio-Rad Laboratories). The validated matrix was applied consistently across all experiments. Thresholds were manually adjusted to lie midway between positive and negative clusters.

### Whole-genome sequencing and coverage analysis

*C. albicans* cells were cultured in 5 mL YPD medium overnight at 30°C with continuous shaking at 200 rpm. The DNA was extracted using the QIAGEN Genomic DNA kit and QIAGEN Genomic-tip20/G, according to the manufacturer’s instructions. Sequencing libraries were prepared by SeqCenter (Pittsburgh, PA) using the Illumina DNA Prep kit and sequenced on an Illumina NovaSeq 6000. The genome sequences and GFF annotation files for the SC5314 reference genome (haplotype A, A22-s07-m01-r130) were downloaded from the *Candida* Genome Database (Braun et al. 2005; Inglis et al. 2012) (http://www.candidagenome.org/). Reads were aligned to the reference genome using Minimap2 v2.17 (Li 2018). Sequence Alignment/Map tools (SAMtools) v1.10 (r783) (Li et al. 2009) and Picard tools v2.23.3 (http://broadinstitute.github.io/picard) were used to filter, sort, and convert the SAM files. Coverage levels were calculated using GATK (McKenna et al. 2010). To estimate chromosome copy numbers, the read alignment depth was calculated for 1 kbp windows across each genome and normalized to the average depth per genome. For published genomes, sequence information is included in **Supplementary Table 1**. Sequence read data for all other genomes have been deposited in the NCBI Sequence Read Archive under BioProject ID PRJNA1354400.

## Results

### Design of a 16-plex chromosome-arm panel using six-color combinations

To quantify the eight *C. albicans* chromosomes, we developed a 16-plex ddPCR assay targeting one locus on each chromosome arm (**Fig. 1A**). The assay employs 6 fluorophores and assigns each chromosome arm/target a unique single- or dual-color code (Santos-Barriopedro 2023) (**Supplementary Table 3**), allowing all 16 loci to be resolved within a single reaction using simple per-channel thresholding. Detection relies on Flex Probe chemistry: target-specific Flex Probes are non-fluorescent oligonucleotides that, upon target amplification, release a universal tag (Tag-oligo) that hybridizes to a fluorophore-labeled Universal Reporter and is extended, generating the fluorescence signal used for droplet calling (**Fig. 1B**).

**Figure 1.**
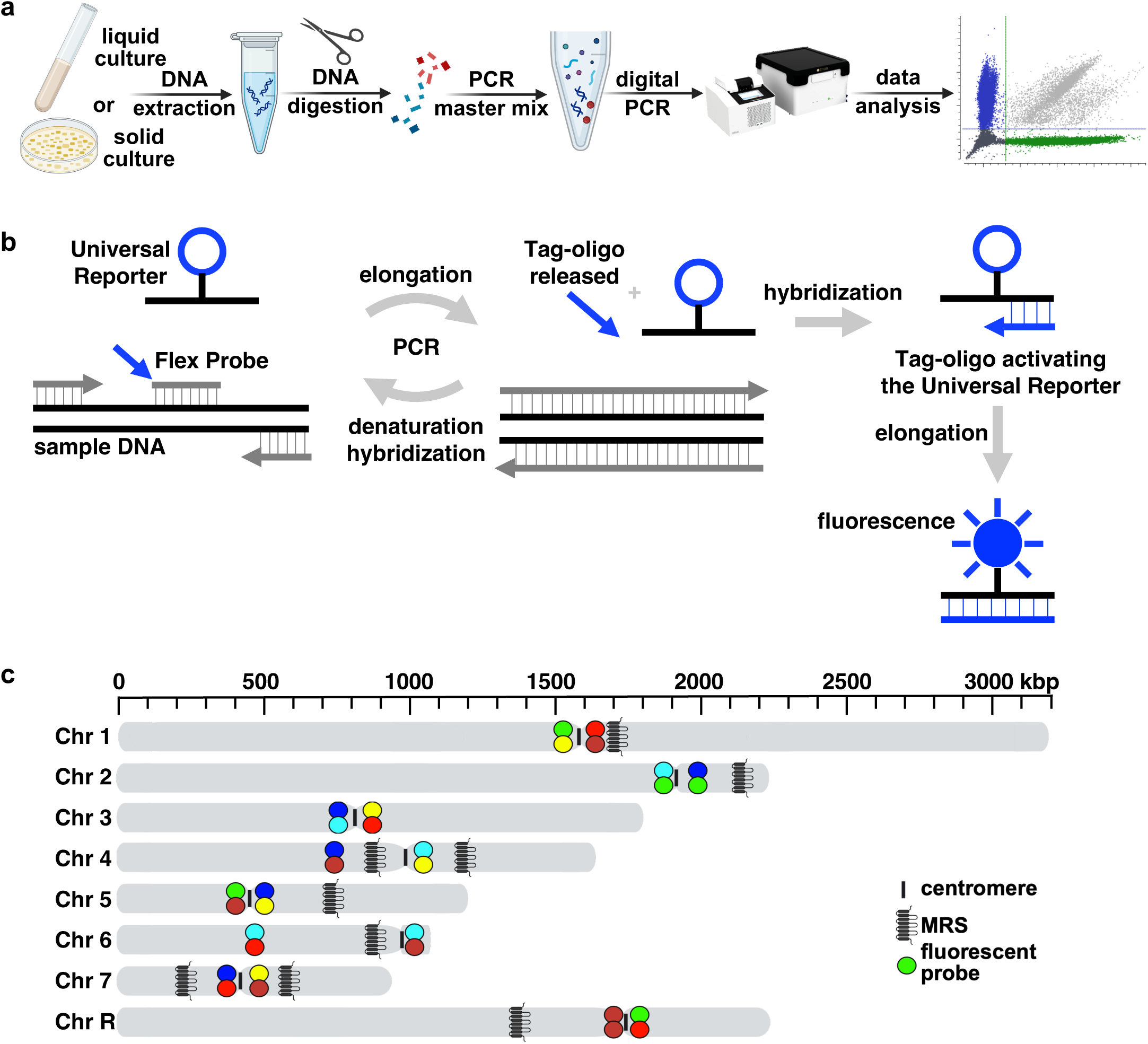
Assay design and fluorescence encoding strategy for 16-plex ddPCR chromosome copy-number profiling in *C. albicans*. (A) Overview of the experimental workflow, including overnight growth of cells (liquid or solid culture), genomic DNA extraction, restriction-enzyme digestion, reaction mix preparation, droplet digital PCR, and downstream data analysis. (B) Flex Probe/Universal Reporter detection chemistry underlying the multiplexing strategy. Forward and reverse primers (gray arrows), Flex probes (gray lines), universal reporters, and genomic DNA are combined in a single reaction. During amplification, target-specific Flex probes are hydrolyzed to release a universal tag (Tag-oligo) that hybridizes to a fluorophore-labeled Universal Reporter. Subsequent elongation generates the target-specific fluorescence signal used for droplet calling, enabling simultaneous discrimination of all 16 loci. (C) Schematic of the eight *C. albicans* chromosomes showing the positions of the 16 targeted loci (one per chromosome arm). Centromeres and major repeat sequence (MRS) regions are indicated. Each arm is assigned a unique single- or dual-color code across six fluorescence channels to enable multiplex detection within a single reaction.

Candidate loci were selected based on normalized WGS coverage near diploid levels at the target site, proximity to the centromere (to avoid telomeric regions prone to mutation and loss of heterozygosity), and association with essential or housekeeping functions (to reduce the likelihood of sequence divergence or structural variation that could impair probe binding). The resulting 16 targets (**Fig. 1C**) provide balanced chromosomal coverage and, because they were chosen from conserved regions, are expected to support consistent probe binding across genetically diverse isolates. As an initial performance check independent of strain background, we tested the panel using equimolar synthetic double-stranded DNA controls (gBlocks) for each chromosome arm. Under these controlled conditions, the measured arm ratios were close to the expected value of 1, supporting the accuracy of the target set and color-coding scheme. Together, this design establishes a scalable, single-reaction framework for genome-wide chromosome copy-number measurement in *C. albicans*.

### Probe redesign and PCR optimization improve multiplex cluster separation

Probes were initially designed as non-competing dual Flex Probe pairs, in which two differently labeled probes bind adjacent sites within the same amplicon (Santos-Barriopedro 2023). Each target was first evaluated individually (simplex) and then combined into the full multiplex panel for single-reaction detection of all 16 loci. When tested on genomic DNA, simplex analysis of two strains (SC5314 (euploid A) and CAY8836-SC (Chr 1,4,5,7 3X)) revealed reproducible deviations from the expected R:L ratio for a subset of chromosomes: Chr1, Chr4, and Chr6 in SC5314, and Chr4 and Chr6 in CAY8836-SC (**Supplementary Fig. 2A**). These deviations persisted in multiplex runs, and an additional imbalance was observed for ChrR in both strains (**Supplementary Fig. 2A**). Inspection of 2D amplitude (dot) plots from simplex reactions localized these effects to probe performance. For SC5314, only one of the two probes designed for each arm produced efficient binding for Chr1, Chr4, and Chr6, whereas in CAY8836-SC, this behavior was restricted to Chr4 and Chr6 (**Supplementary Fig. 2B**).

Because each arm should contribute ∼50% of the double-positive signal, inefficient detection of one arm depletes the double-positive cluster and inflates single-color droplets, yielding an apparent arm imbalance. Notably, ChrRL is the only target encoded in a single channel (IR), and Chr6R generated a marked excess of IR-only droplets (**Supplementary Table 5**). Thus, a fraction of Chr6R molecules that should have contributed to the double-positive population were instead detected only in IR, biasing the inferred ChrR R:L ratio. We hypothesized that single-nucleotide polymorphisms (SNPs) and/or local DNA conformation impaired probe hybridization. To correct these probe-specific biases, we redesigned the probes for Chr1L, Chr4R, and Chr6R using a competing dual Flex Probe configuration, in which both probes target the same region and compete for hybridization (Santos-Barriopedro 2023). For each target, the redesigned pair was based on the better-performing sequence from the original non-competing design.

Additional optimization steps included increasing the probe concentration from 0.5 µM to 1 µM and extending the final elongation step from 10 to 15 min. These adjustments increased signal amplitude and reduced stochastic detection variability, particularly under multiplex conditions. Consistent with this, average separability scores (indicating how well the positive and negative distributions are separated, **Supplementary Fig. 1**) improved after probe redesign and reaction optimization, yielding well-resolved positive and negative droplet populations across all targets. The longer final elongation improved separability (**Supplementary Fig. 3A**), likely by allowing additional time for the Tag-oligo hybridized to the Universal Reporter to elongate and release the fluorophore. Because the optimal elongation time is assay-dependent, it can be adjusted to improve amplification efficacy and maximize cluster height, although overly long elongation steps may occasionally produce low-level positive droplets in the no-template control.

Together, these refinements produced distinct, non-overlapping signal clusters for all 16 loci within a single ddPCR reaction. Partition clustering analysis reliably separated positive and negative droplets, enabling direct absolute quantification of each target and robust genome-wide aneuploidy measurement in *C. albicans*.

### Spillover compensation improves multiplex droplet calling

Droplet calling and quantification were performed in Crystal Miner. Because the naica system detects six fluorophores across six channels, partial spectral overlap (“spillover”) can distort droplet amplitudes and bias multiplex quantification. Although each fluorophore is primarily excited and detected in its own channel, low-level excitation by other light sources and broad emission spectra can introduce measurable signal into non-primary channels. To correct this, we generated an assay-specific spillover compensation matrix using chambers containing single-color probe sets. For each fluorophore, we quantified the fraction of its signal detected in the other channels and used these values to build a spillover compensation matrix. The matrix was then inverted and applied to raw fluorescence measurements to obtain spillover-corrected fluorophore intensities and adjusted for background fluorescence. This correction yielded cleaner one-dimensional amplitude distributions for thresholding (**Supplementary Fig. 3B**) and restored the orthogonal cluster geometry in 2D amplitude plots, improving separation between negative and positive droplet populations (**Supplementary Fig. 3C**). The resulting matrix can be saved and reused for all runs performed with this assay. After compensation, thresholds separating positive and negative droplets were manually set in each channel, and the population editor was used to define single- and dual-color codes for positive droplets of each target (e.g., *CCT2* in the blue and green channels and *SSK2* in the teal and yellow channels, **Supplementary Fig. 3C**). Using Poisson statistics, the software then estimated target concentrations and reported confidence intervals. Together, spillover correction and consistent droplet calling across runs improve high-order multiplex quantification.

### Column-based DNA extraction and higher DNA input improve 16-plex copy number accuracy

To assess how genomic DNA preparation affects copy-number calling, we compared a widely used phenol–chloroform protocol with a commercial column-based kit (QIAGEN) across five *C. albicans* isolates spanning euploid (SC5314), simple aneuploid (Chr3 1X, ChrR 3X), and complex aneuploid (Chr3,6 3X, Chr6,R 4X) karyotypes. Baseline chromosome copy numbers were first established by Illumina WGS (**Fig. 2A**). When these samples were analyzed with the 16-plex ddPCR assay, phenol–chloroform DNA produced noisier copy-number profiles and frequent deviations from the WGS karyotype (**Fig. 2B**). In contrast, column-purified DNA yielded chromosome copy-number estimates that closely matched WGS across all isolates (**Fig. 2B**), reduced technical variance (from a mean SD of 0.71 to 0.38), and increased the fraction of positive droplets per reaction (from a total of 3.25% to 21.21%) (**Supplementary Fig. 4**).

**Figure 2.**
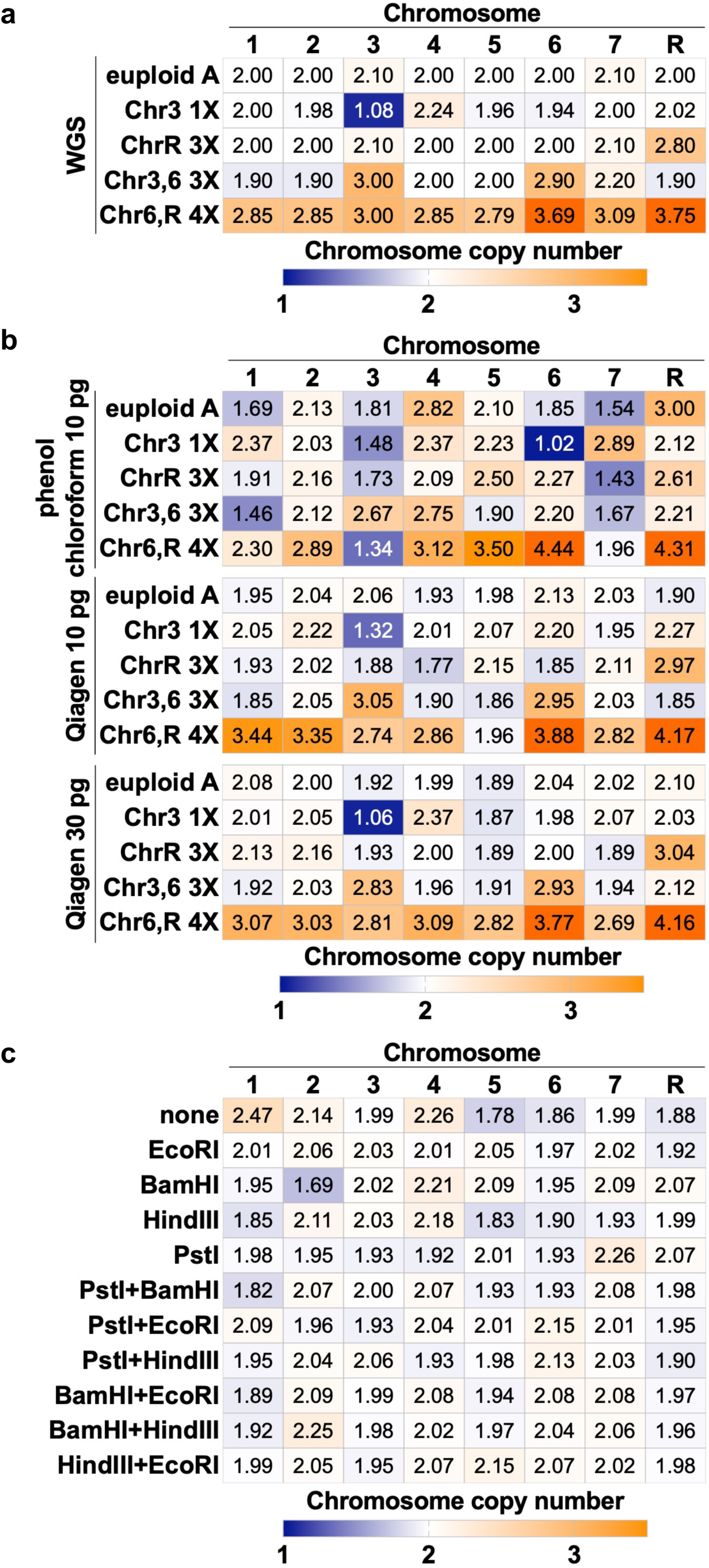
Genomic DNA preparation steps that improve 16-plex ddPCR accuracy. (A) Whole-genome sequencing-derived chromosome copy-number profiles for a panel comprising one euploid reference isolate and four aneuploid isolates representing simple and complex karyotypes. Values are shown as chromosome-level copy numbers. (B) Heatmaps of ddPCR-derived chromosome copy-number estimates for the same isolates under different extraction/input conditions, comparing phenol–chloroform extraction (10 pg/µL input) to column-based purification (QIAGEN; 10 pg/µL and 30 pg/µL input). (C) Effect of restriction-enzyme digestion on ddPCR accuracy using the euploid reference strain SC5314 (euploid A). Heatmaps show chromosome copy-number estimates obtained without digestion (none) or after digestion with individual enzymes or enzyme combinations.

We next asked whether increasing DNA input could further improve assay precision. While 10 pg/µL of column-purified DNA produced acceptable profiles, raising the input to 30 pg/µL increased concordance with WGS (**Fig. 2B**), improved reproducibility across replicates (from a mean SD of 0.38 to 0.16), and further increased the positive droplet fraction from 21.21% to 38.7% (**Supplementary Fig. 4**). Based on these results, the optimized workflow uses 30 pg of column-purified gDNA per 5 µL reaction to maximize target detection and minimize false aneuploid calls.

### DNA digestion increases droplet resolution and copy number accuracy in the 16-plex assay

High-molecular-weight genomic DNA can interfere with droplet formation and reduce amplification uniformity across targets. To test whether template fragmentation enhances assay performance, genomic DNA from the euploid reference strain SC5314 (euploid A) was digested with EcoRI, BamHI, HindIII, or PstI, either individually or in pairwise combinations, before amplification. Enzymes were selected to avoid predicted cut sites within the 16 amplicons (**Supplementary Table 2**).

Across conditions, digestion improved the accuracy of chromosome copy-number estimates relative to undigested DNA (**Fig. 2C**). While single-enzyme digests provided modest gains, enzyme combinations produced the most consistent improvements, particularly when PstI was combined with either EcoRI or HindIII. This was reflected by reduced variability between replicates in copy-number estimates and higher separability scores across chromosome arms (**Supplementary Fig. 5A-B**). Digestion also increased the fraction of positive droplets (**Supplementary Fig. 5C**), consistent with improved accessibility of amplifiable templates and reduced stochastic dropout. Undigested DNA tended to overestimate copy number for specific chromosomes, particularly for Chr1 and Chr4, whereas digestion corrected these biases and returned copy numbers closer to the expected disomic baseline (**Fig. 2C** and **Supplementary Fig. 5D**). Taken together, restriction digestion improved both droplet cluster resolution and quantitative accuracy. Consequently, we incorporated DNA pre-digestion with PstI+HindIII as a standard step in the optimized 16-plex workflow.

### The 16-plex assay sensitively detects mosaic aneuploidy and matches WGS across diverse karyotypes

To test sensitivity for detecting emerging aneuploid subpopulations, we generated defined mixtures of disomic and trisomic cultures from the same strain background (ChrR 2X:3X) at 25%, 50%, and 75% trisomic fractions. The assay revealed a progressive increase in ChrR copy number that closely matched the expected mixture averages, and reliably distinguished trisomy when only 25% of the population carried the extra chromosome (**Fig. 3A**). The same trend was observed at the chromosome-arm level for RL and RR (**Supplementary Table 6**), indicating that the 16-plex format can resolve mosaic aneuploidy at low abundance despite the sampling noise inherent to droplet partitioning.

**Figure 3.**
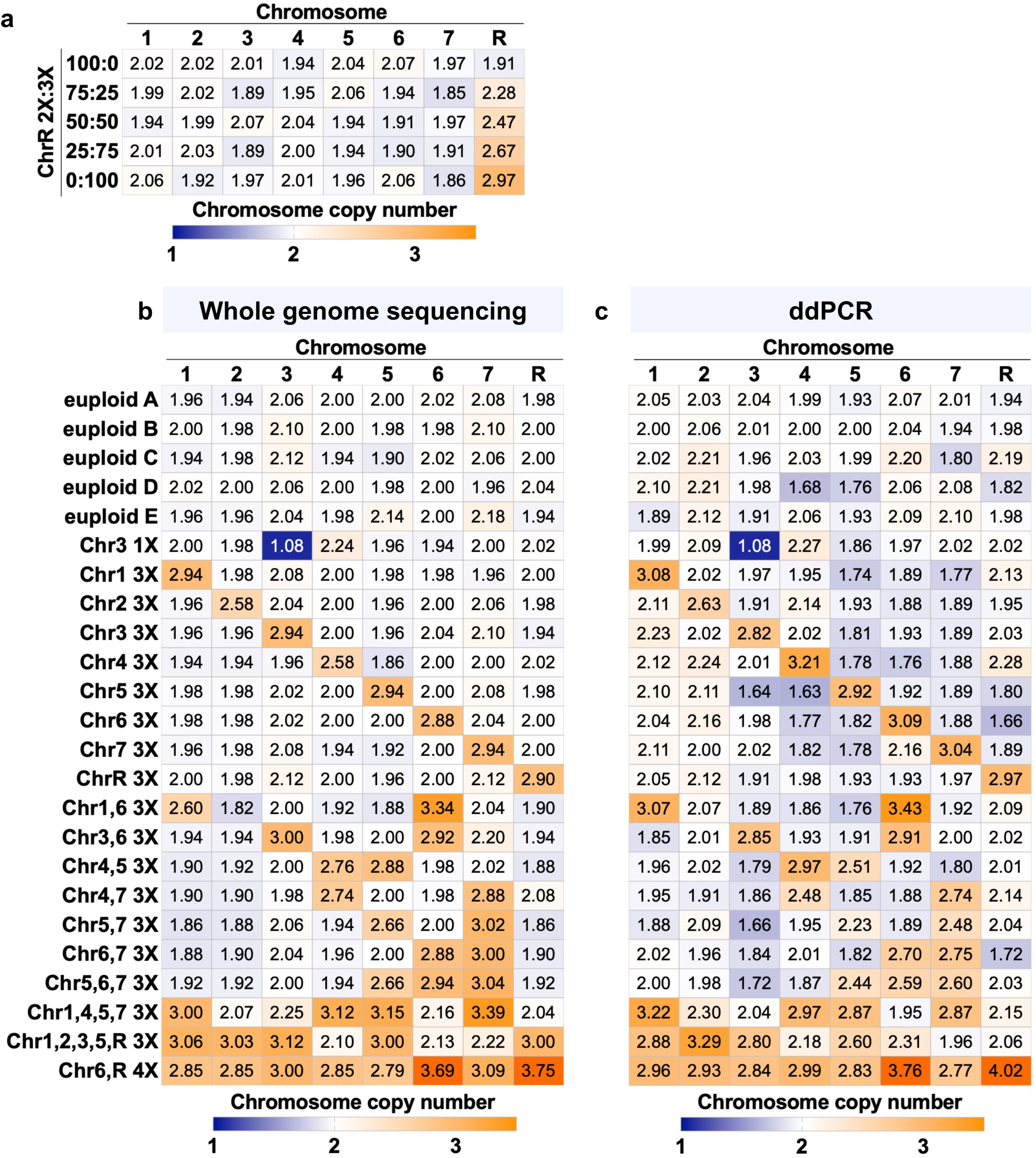
Assay performance across mixed populations and diverse isolate karyotypes. (A) Detection of mosaic aneuploidy using defined mixtures of disomic (euploid B, ChrR 2X) and aneuploid (ChrR 3X) subpopulations. (B) WGS-based chromosome copy-number profiles for 24 isolates spanning euploid, single-aneuploid, and complex karyotypes. (C) Corresponding ddPCR-based chromosome copy-number estimates for the same isolate panel.

We next assessed species specificity using genomic DNA from *C. dubliniensis* and *C. tropicalis*. Under standard cycling conditions, none of the 16 targets generated positive droplets, supporting assay specificity for *C. albicans* and suggesting that detection of related species would require redesigned primers/probes.

Finally, we benchmarked ddPCR copy-number estimates against Illumina WGS across a panel spanning euploid isolates and increasingly complex aneuploid karyotypes (**Fig. 3B**). Euploid controls clustered tightly around the expected disomic baseline, and the assay correctly identified single-chromosome aneuploidies across all eight chromosomes, including Chr3 monosomy and distinct trisomies (**Fig. 3C**). Isolates carrying two supernumerary chromosomes were also called accurately, with strong agreement between ddPCR and WGS (**Fig. 3B-C**) and consistent left/right arm estimates (**Supplementary Table 7**). In highly aneuploid isolates (≥4 altered chromosomes), ddPCR still captured broad copy-number elevations and flagged complex karyotypes, but complete resolution became less consistent, occasionally underestimating one affected chromosome, likely because calling becomes more challenging when many targets shift at once in highly aneuploid genomes (**Fig. 3C**).

Overall, these results show that the 16-plex ddPCR assay provides WGS-concordant copy-number calls for euploid, single-, and dual-aneuploid karyotypes, while remaining sensitive enough to detect low-frequency trisomic subclones and to flag isolates with more complex chromosomal alterations.

## Discussion

Here, we developed and validated a 16-plex ddPCR assay for genome-wide chromosome copy-number measurement in *C. albicans.* The assay combines digital quantification with high-order multiplexing through color-combination encoding across six fluorescence channels, enabling simultaneous readout of all 16 chromosome arms in a single reaction. Targeting both chromosome arms gives a built-in cross-check (left vs. right) and helps capture large arm-scale copy-number changes that single-locus qPCR assays can miss. In practice, this workflow provides a rapid alternative to sequencing: from gDNA to copy-number calls can be completed within a day, without library preparation or bioinformatics.

As with any targeted assay, performance depends on the complementarity between probe and template. Locus-specific polymorphisms or small indels in primer/probe binding sites can reduce hybridization efficiency and bias local quantification in genetically divergent isolates. In addition, the resolution is limited to the assayed loci: partial aneuploidies that do not include the selected target region will not be detected, and breakpoints cannot be mapped. Finally, high-order multiplexing requires stable fluorescence calibration. Spectral overlap and variability between replicates can affect cluster positions if compensation and thresholding are not standardized. These limitations can be mitigated by (i) selecting conserved targets (as done here) and redesigning individual probe sets when strain-specific dropout is observed, (ii) including a reference strain with a known karyotype, and (iii) using fixed instrument templates and a fixed spillover compensation matrix to stabilize cluster geometry across experiments.

Across a diverse isolate panel, the 16-plex assay showed strong agreement with WGS for euploid strains and for isolates carrying 1-3 supernumerary chromosomes, and it sensitively detected mosaic trisomy down to 25% abundance in mixed populations. Performance was less consistent in highly aneuploid genomes, where many chromosomes deviate simultaneously and accurate calling becomes more challenging. Nevertheless, even in these cases, the assay reliably flagged complex karyotypes and captured broad patterns of copy-number variation. Together, these results define a practical operating range: highly accurate for simple aneuploid states, and informative for more complex karyotypes where follow-up by WGS may be warranted.

Relative to WGS, ddPCR offers advantages in speed, cost, and sample requirements. It is best suited for large-scale screening experiments where many isolates must be profiled quickly, as well as for settings where sequencing infrastructure is limited. The assay’s reliance on droplet partitioning also makes it compatible with low DNA inputs and heterogeneous populations, making it relevant for clinical isolates and samples where aneuploid subpopulations may be emerging. Our optimization results underscore, however, that standardized sample preparation is essential: column-based extraction, higher DNA input, and restriction digestion substantially improved positive droplet fraction, cluster separation, and copy-number accuracy.

A key strength of the assay is its modularity. Because multiplex structure is encoded by probe sequences and fluorophore combinations, panels can be reconfigured to address new biological questions without changing the underlying workflow. For example, the same design principles could be extended to track recurrent aneuploidies linked to drug resistance or adaptation, to include loci reporting focal amplifications, or to incorporate species- or clade-informative markers alongside copy-number targets. More broadly, by redesigning primer/probe sequences while preserving the color-combination logic, this approach should translate to other fungal species where copy number variation contributes to stress resistance and host adaptation.

Overall, this 16-plex ddPCR assay bridges a practical gap between comprehensive but resource-intensive sequencing and low-multiplex assays. It provides an accessible platform for quantitative monitoring of genome instability, enabling rapid, scalable readout of karyotypic variation in both experimental and clinical contexts.

## Data availability

Sequence read data generated in this study have been deposited in the NCBI Sequence Read Archive under BioProject PRJNA1354400. Accession information for previously published genomes analyzed here is provided in **Supplementary Table 1**. Strains are available upon request. The authors affirm that all data necessary for confirming the conclusions of the article are present within the article, figures, and tables.

## Acknowledgements

We thank members of the Fungal Heterogeneity Group for helpful discussions.

## Supplementary Figure Legends

**Supplementary Figure 1.**
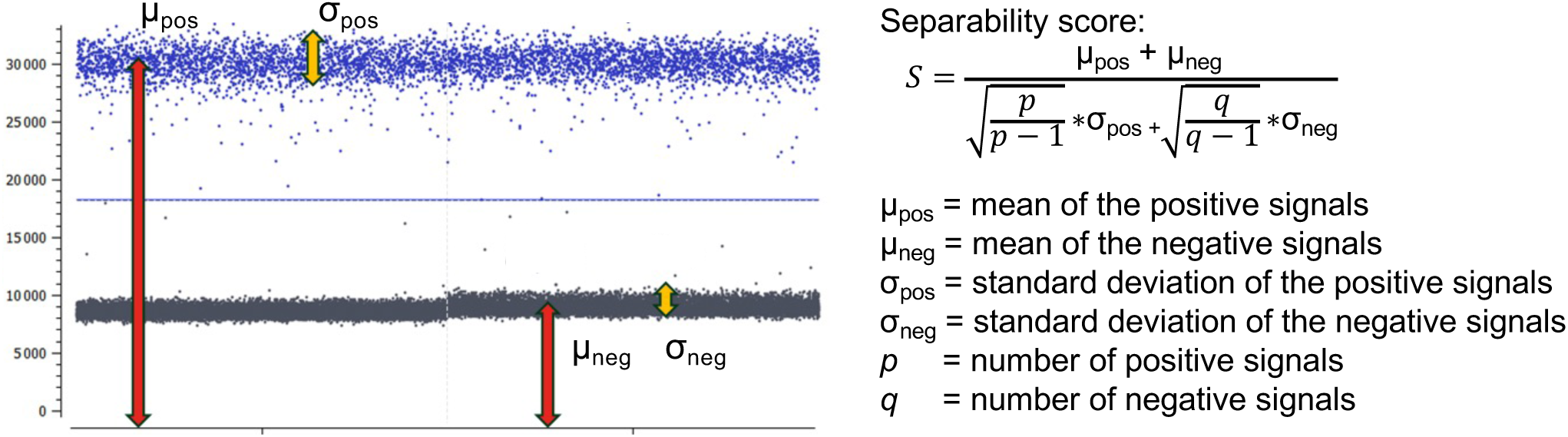
Separability score used to quantify droplet-cluster resolution. Definition of the separability score (S) computed from the mean and dispersion of positive and negative droplet amplitude distributions and the number of droplets in each population. Higher values indicate better separation between positive and negative droplets and improved robustness of thresholding.

**Supplementary Figure 2.**
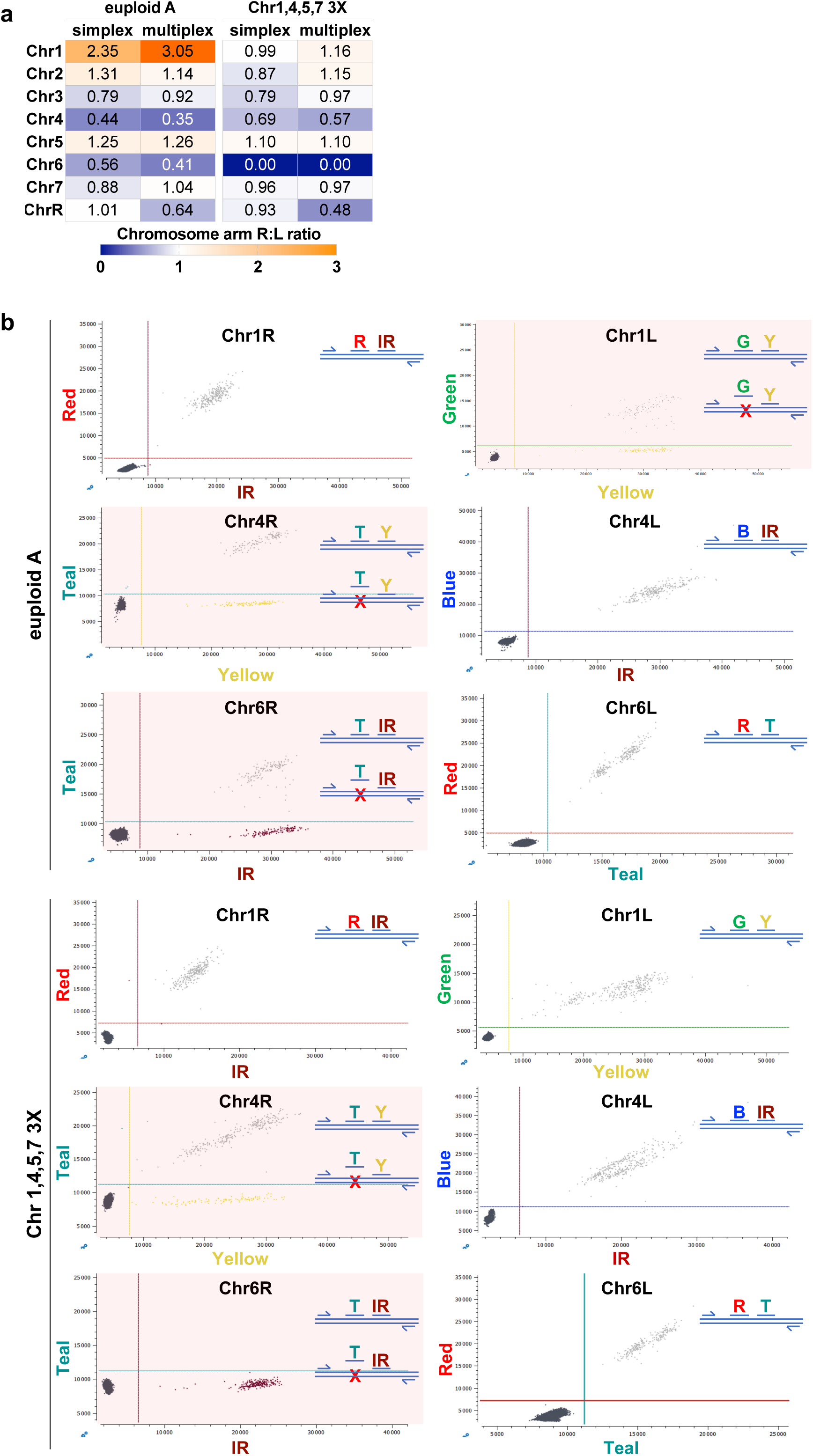
Strain-dependent chromosome arm imbalances reveal probe-specific underperformance. (A) Heatmap of chromosome arm R:L ratios for isolates SC5314 (euploid A) and CAY8836-SC (Chr1,4,5,7, 3X) measured in simplex and multiplex. For each chromosome, the ratio was calculated as the copy-number estimate for the right-arm (R) target divided by the left-arm (L) target; values near 1 indicate balanced detection of both arms, whereas deviations reflect arm-specific bias (notably for Chr1, Chr4, Chr6, and ChrR). (B) Representative 2D amplitude (dot) plots from simplex reactions for arm targets showing the underlying probe behavior in euploid A (top) and Chr1,4,5,7, 3X (bottom) isolates. For Chr1L, Chr4R, and Chr6R, one probe in the original non-competing dual-probe pair yields inefficient detection (shown by the crossed-out probe), reducing the expected double-positive population and inflating single-color droplets. In contrast, the corresponding partner-arm targets (Chr1R, Chr4L, Chr6L) show the expected well-separated positive clusters in their assigned channel pairs.

**Supplementary Figure 3.**
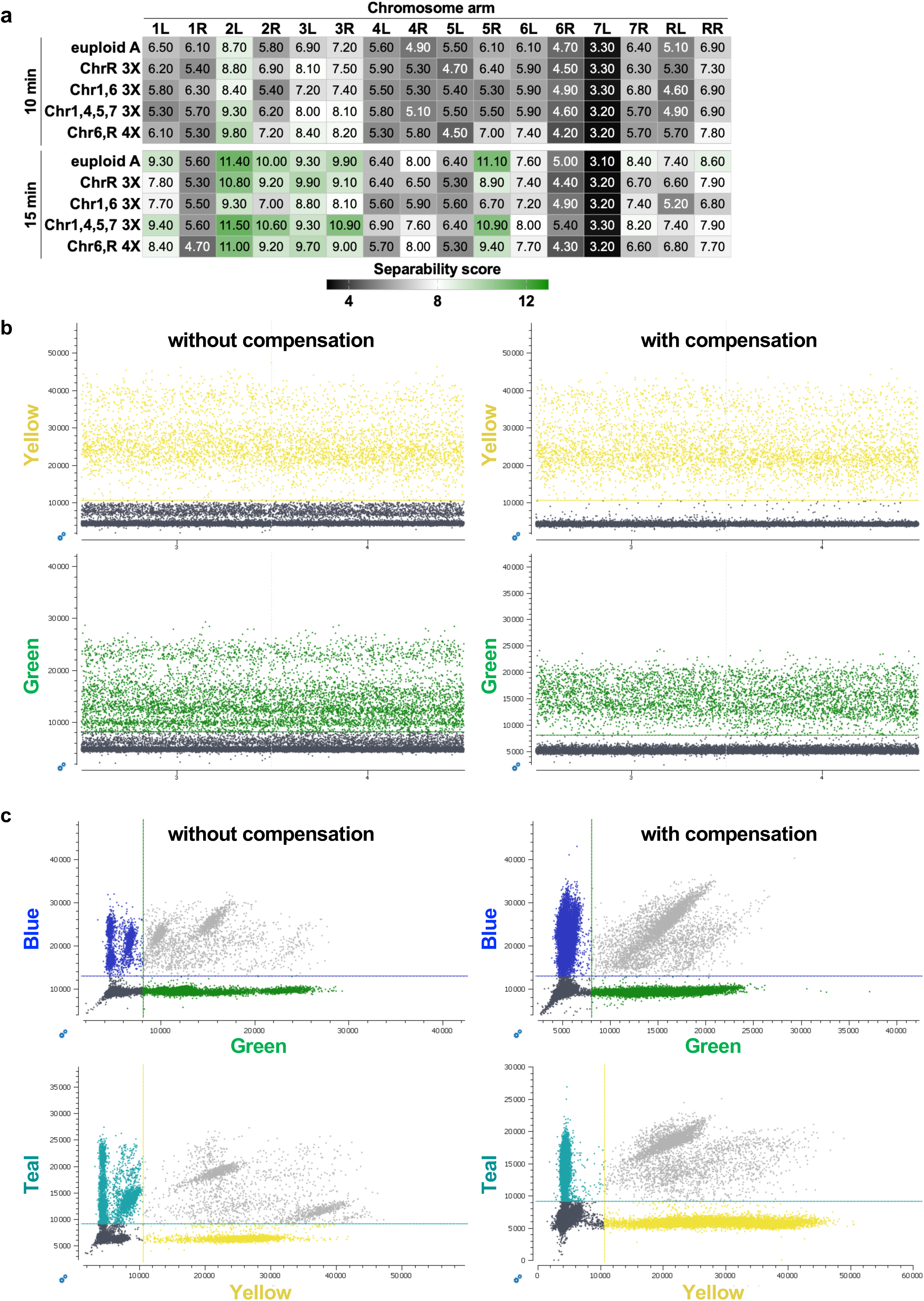
Final elongation time and spillover compensation improve cluster separation in multiplex reactions. (A) Heatmap of separability scores for each chromosome-arm target (columns) across representative isolates, comparing a 10 min versus 15 min final elongation step. Higher scores indicate better separation between positive and negative droplet populations. (B) Representative 1D amplitude plots in the yellow and green channels before and after applying spillover compensation. Compensation yields a cleaner separation of positive (colored) and negative (gray) droplet populations. (C) Representative 2D amplitude (dot) plots (Blue vs Green; Teal vs Yellow) before and after compensation. Spillover correction restores the expected cluster geometry and increases separation between negative (gray) and positive (colored) droplet populations, supporting more reliable droplet calling.

**Supplementary Figure 4.**
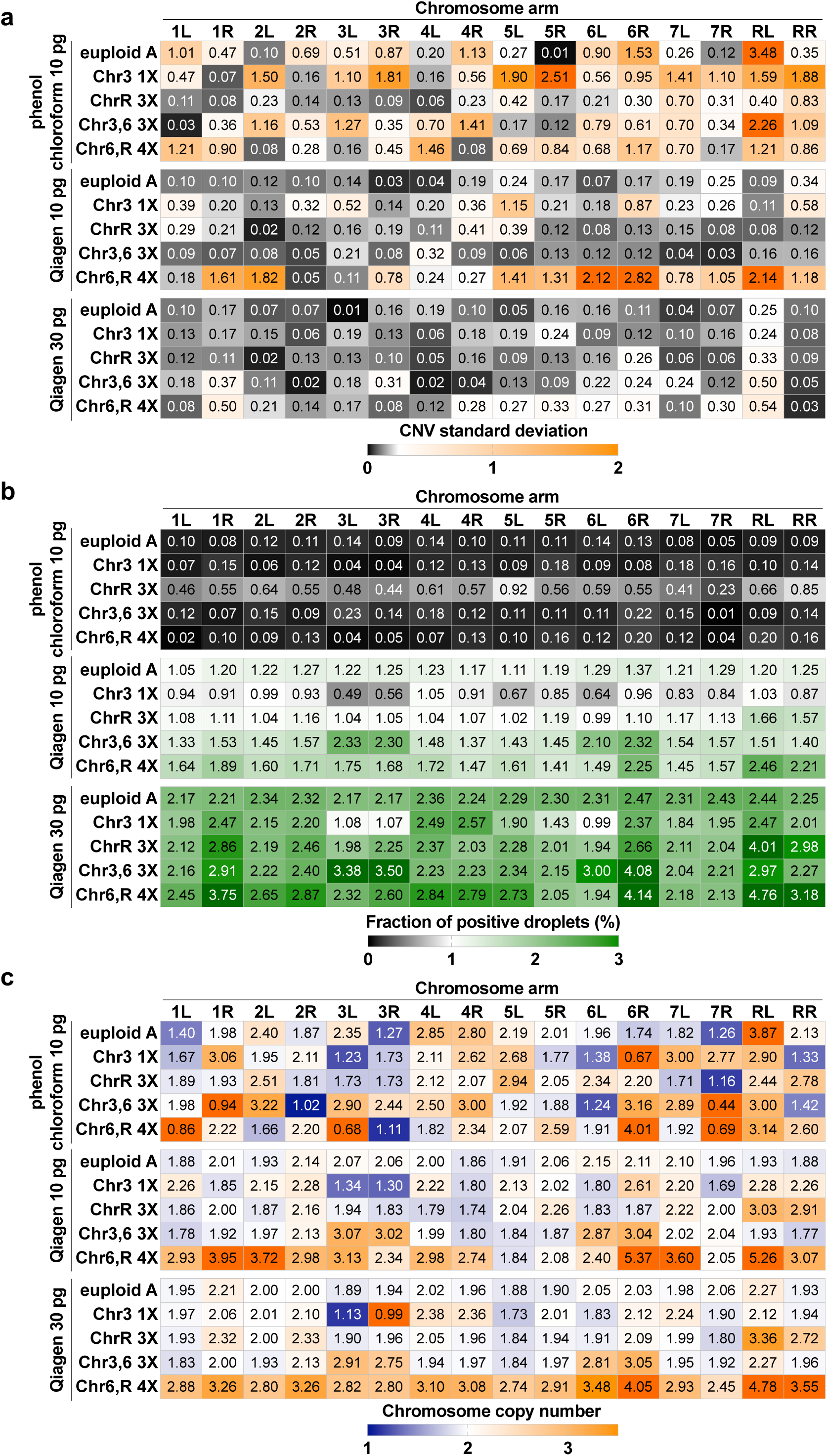
Effects of DNA extraction method and DNA input amount on the performance of the 16-plex assay across euploid and aneuploid isolates. (A) Standard deviation of chromosome copy-number estimates across biological replicates as a measure of technical variance. (B) Fraction of positive droplets per reaction/condition, reflecting effective amplifiable template availability. (C) Chromosome copy-number estimates under each condition, highlighting false-positive calls under suboptimal preparation.

**Supplementary Figure 5.**
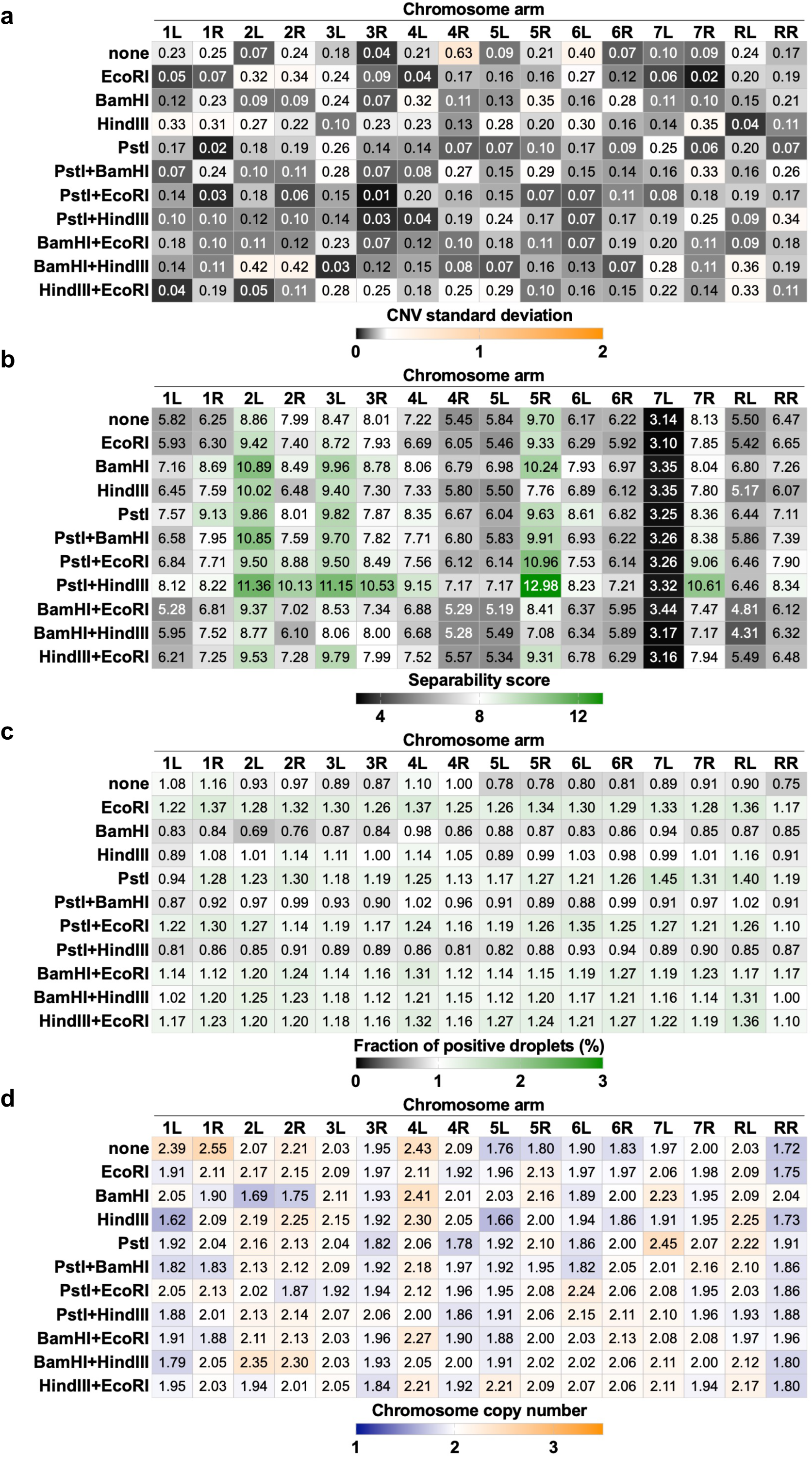
Restriction-enzyme digestion improves droplet resolution and assay consistency. Performance metrics for undigested DNA versus digestion with single enzymes or enzyme combinations. Heatmaps summarize changes following SC5314 (euploid A) DNA digestion in (A) variability of chromosome copy-number estimates (SD across replicates), (B) droplet separability scores, (C) fraction of positive droplets, and (D) chromosome copy-number profiles.

## Supplementary Table Legends

**Supplementary Table 1**. *Candida* isolates used in this study. List of isolates analyzed, including strain names and aliases, sequence information, and literature references where applicable.

**Supplementary Table 2.** Panel of restriction enzymes evaluated for genomic DNA digestion and their predicted cleavage within the amplified target region of each chromosome arm.

**Supplementary Table 3.** Target loci and fluorophore combinations used in the 16-plex ddPCR assay. Each *C. albicans* chromosome arm is represented by a single target gene detected through a unique fluorophore or pair of fluorophores across six color channels. The combination of fluorophores defines a distinct fluorescence signature for each target. The final column indicates whether the two probes for that target were designed with competitive or non-competitive binding.

**Supplementary Table 4.** Primers and Flex probes used in this study.

**Supplementary Table 5.** Percentage of target detection for each chromosome arm for strains SC5314 (euploid A) and CAY8836-SC (Chr 1,4,5,7 3X), calculated by dividing the number of positive droplets per arm by the total number of droplets for each chromosome.

**Supplementary Table 6.** Chromosome-arm copy number estimates in defined mixed populations of euploid B (ChrR disomic, 2X) and ChrR 3X (ChrR trisomic, 3X) isolates from the same genetic background (CAY6440) at 0%, 25%, 50%, 75%, and 100% trisomic fractions.

**Supplementary Table 7.** Chromosome-arm copy number estimates across a panel of 24 isolates, showing concordance between left and right arms for most chromosomes and highlighting isolates where arm estimates diverge, consistent with arm-biased copy-number changes or structural variation spanning one target.

